# N-Acetyl-l-Leucine (NALL) rescues inter-organelle communication in Niemann-Pick disease type-C patient cells

**DOI:** 10.1101/2025.04.27.650877

**Authors:** Szilvia Kiraly, Andrea Martello, Frances M. Platt, Emily R. Eden

## Abstract

Niemann-Pick disease type-C (NPC) is a progressive neurodegenerative disease caused by loss-of-function mutations in *NPC1* or *NPC2*. In NPC patient cells lacking functional NPC proteins, lipids accumulate in lysosomes causing severe lysosomal storage disease. Surprisingly, lipid accumulation caused by defects in the lysosomal membrane protein NPC1 is strongly associated with mitochondrial dysfunction. The mechanism of this coupled dysfunction is not fully understood, but a recently approved NPC therapeutic, N-Acetyl-l-Leucine (NALL), reverses both lysosomal and mitochondrial phenotypes in NPC patient cells. Our data indicate that direct inter-organelle communication through lysosome membrane contact sites with mitochondria contribute to the coupled organelle dysfunction in NPC. We find that mitochondria:lysosome contact sites are expanded in NPC, dependent on accumulation of lysosomal cholesterol and that NALL rescues the aberrant contact sites. We further identify a direct correlation between mitochondria:lysosome contact site expansion and mitochondrial dysfunction and propose that normalisation of contacts sites contributes to the coupled restoration of lysosome and mitochondrial function by NALL. We further find that NALL-mediated normalisation of lysosomal contact sites also correlates with restoration of autophagic flux and lysosome repair in NPC patient cells.

## Introduction

Niemann-Pick disease type-C (NPC) is a progressive neurodegenerative lysosomal storage disorder most typically presenting in infancy/childhood. NPC is caused by loss of function mutations in one of two lipid transport proteins, NPC1 and NPC2, that together facilitate the egress of dietary cholesterol from the endocytic pathway. NPC1 is a large multipass transmembrane domain protein that can transport luminal cholesterol and sphingosine (1) across the glycocalyx to the limiting membrane of LE/Lys through a hydrophobic tunnel (2). In contrast, NPC2 is a small luminal protein that delivers cholesterol from intraluminal membranes to NPC1 for transport to the limiting membrane (3). In the absence of functional NPC1 or NPC2 proteins, cholesterol and sphingolipids accumulate in late endosomes and lysosomes (LE/Lys), which become dysfunctional (4). Curiously, mitochondrial dysfunction is also associated with NPC (5) and is a common pathogenic feature of lysosomal storage and neurodegenerative disease (6), suggesting a key role of mitochondria in NPC disease pathogenesis. Why the failure to move lipids from the lysosome to the ER in NPC disease is associated with mitochondrial dysfunction is not yet fully understood, but one potential mechanism is transport of excess cholesterol from lipid-laden lysosomes to mitochondria. In NPC1-deficient cells, cholesterol was found not only to accumulate in lysosomes, but also in mitochondria (7, 8), with mitochondrial sterol accumulation mediated by the LE/Lys sterol transfer protein STARD3 (8). Mitochondrial cholesterol accumulation is strongly implicated in oxidative stress and changes in metabolic homeostasis (9, 10) and is therefore likely to contribute to mitochondrial dysfunction in NPC.

Both lysosomes and mitochondria are essential for regulating cellular metabolism and complex inter-organelle crosstalk coordinates their metabolic roles (11). During the last decade we have begun to appreciate the importance of mitochondria:LE/Lys contact sites (MLCs) in inter-organellar communication. Membrane contact sites are highly regulated domains where neighbouring organelles are in very close apposition (typically 5-40 nm apart). We have previously identified a role for NPC1, in addition to it’s well established role in transporting cholesterol from the lumen to the limiting membrane of LE/Lys, in stabilising LE/Lys contact sites with the ER through interaction with ER-resident lipid transfer protein Gramd1b (12). Thus in NPC1-deficent cells ER:LE/Lys contact sites were reduced, but in contrast, we found a striking expansion of mitochondria:LE/Lys contact that, like mitochondrial cholesterol accumulation in NPC, is also dependent on STARD3 (13). Since their identification in yeast approximately a decade ago, MLCs have been implicated in regulating far-reaching processes important for the maintenance of cellular homeostasis (14). Although expanded MLCs have been implicated in coupled lysosome:mitochondria dysfunction in other disease settings, especially Parkinson’s disease (15), MLC regulation and their role in lysosome-mitochondrial crosstalk is poorly understood.

Here we investigate the effect of N-Acetyl-l-Leucine (NALL) on coupled lysosome:mitochondria dysfunction in NPC patient-derived fibroblasts. NALL is a modified derivative of the branched-chain essential amino acid leucine that has been shown to significantly reduce NPC disease progression (16) and was FDA approved as an NPC therapeutic in 2024. Acetylation converts leucine into an anion that becomes a substrate for monocarboxylate transporters (MCT), bypassing the rate-limiting uptake by the low capacity L-type amino acid transporter (LAT1) that is responsible for non-acetylated leucine uptake (17). NALL was found to rescue both lysosomal lipid accumulation and mitochondrial respiration in cellular and animal models of NPC (18), with rescue of autophagic flux proposed as a potential therapeutic mechanism (19).

In this study, we find that NALL-driven rescue of lysosomal and mitochondrial phenotypes in NPC1 patient cells correlates with a reduction in the extent of MLCs and increased LE/Lys contact with the ER. Clearance of cholesterol from LE/Lys by NALL treatment is associated with increased esterification and lipid droplet formation, recovery of lysosome repair and restoration of autophagic flux. Thus normalisation of LE/Lys contact sites with the ER and mitochondria following treatment with NALL correlates with coupled rescue of lysosome and mitochondrial dysfunction and restoration of metabolic homeostasis.

## Results

### Rescue of lysosome expansion and lipid accumulation in NPC patient cells by NALL treatment

The primary fibroblasts used in this study, described in Table-1, are from two unrelated healthy control donors (HC1, HC2) and three unrelated NPC patients (three with NPC1 mutations NPC1-P1, -P2 and -P3). Quantitation of filipin-stained cholesterol revealed a time-dependent effect of NALL treatment on lysosomal cholesterol accumulation in NPC patient-derived cells (Supplemental Figure S1A and S1B). Prior to NALL treatment, filipin-stained cholesterol was significantly increased in all NPC patient cells compared with controls, but following a 72 h incubation with NALL, there was no longer any significant difference between NPC and control cells (Figure 1A and 1B/Supplemental Figure S1C and S1D).

**Figure 1.**
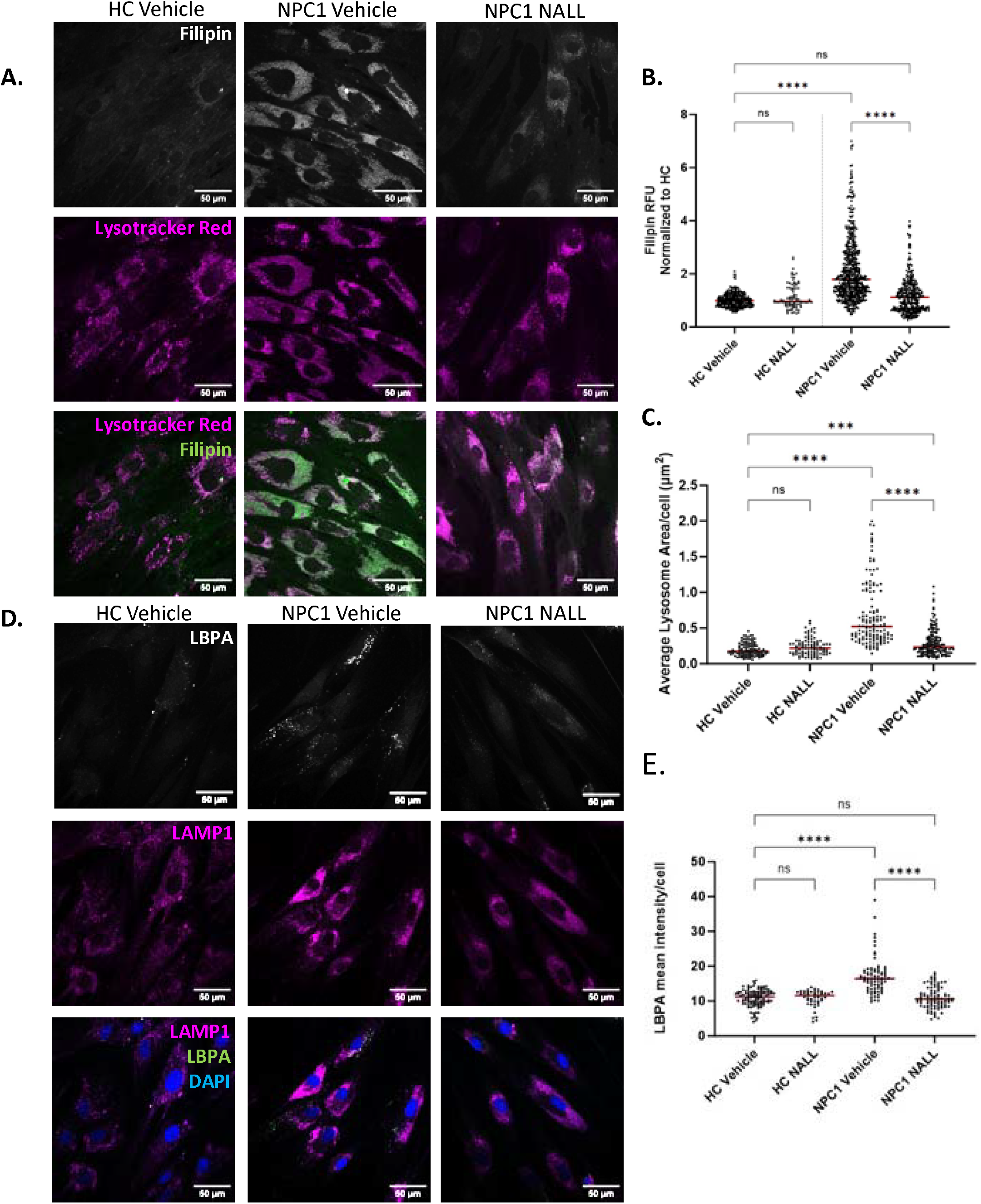
NALL rescues LE/Lys lipid accumulation and expansion. Primary fibroblasts from healthy control donors (HC) or NPC1 patients were treated with DMSO (vehicle) or 1mM NALL (NALL) for 72 hours followed by 30 min with lysotracker. Cells were fixed prior to imaging of filipin-stained cholesterol (A, quantitation in B), lysotracker (A, quantitation in C) or anti-LBPA staining (D, quantitation in E). Scale bar, 50μm. Scatter plots show pooled data from 3 NPC patient cell lines and normalized to the healthy control average, centre lines show medians (B HC: *n* = 308, HC NALL: n= 92 NPC1: *n* = 572, NPC1 NALL: *n* = 340 cells C) HC: *n* = 301, HC NALL: n= 284 NPC1: *n* = 492, NPC1 NALL: *n* = 648 organelles E) HC: *n* = 78, HC NALL: n=74 NPC1: *n* = 172, NPC1 NALL: *n* = 204 cells). Data analysed by one-way ANOVA followed by Tukey’s multiple comparisons test comparing all groups (* p ≤ 0.05, ** p ≤ 0.01, *** p ≤ 0.001, ns = non-

Lipid accumulation in NPC is associated with LE/Lys expansion and LE/Lys volume can be useful both as a biomarker of lysosomal storage diseases and an indicator of response to therapeutics (20). To measure LE/Lys expansion, cells were cultured with LysoTracker, which revealed an increase in LE/lysosome area in NPC1 patient cells compared with controls that was reversed by treatment with NALL (Figure 1A and 1C).

The atypical phospholipid lysobisphosphatidic acid (LBPA), also known as bis(monoacylglycerol)phosphate (BMP), localises to intraluminal membranes of a subpopulation of LE/Lys that do not contain EGF receptor (21). Cholesterol accumulates within the LBPA-positive population of LE/Lys in NPC and LBPA is increased in NPC1-inhibited cells (22). Counterintuitively, further increasing LBPA has been proposed as a potential NPC therapeutic strategy since addition of its precursor PG, which increases LBPA in LE/Lys, results in an NPC2-dependent reduction in cholesterol accumulation in NPC (23), thought to be mediated by the induction of autophagy (24). Consistent with previous reports, we found increased LBPA in in LE/Lys of NPC cells compared with controls. LBPA accumulation in NPC cells was reversed following a 72h treatment with NALL (Figure 1D and 1E). These data indicate that NALL rescues LE/Lys phenotypes in NPC, though the mechanism remains unclear.

### NALL increases ER contact with LE/Lys, cholesterol esterification and lipid droplet formation

We next examined the effects of NALL treatment on LE/Lys at the ultrastructural level by electron microscopy (EM). Whereas LE/Lys circularity is reduced, and density is increased in NPC patient cells, there was no significant difference between these parameters in NPC and control cells following NALL treatment 72 hours (Figure 2A and 2B). However, although LE/Lys shape was normalised, there was a higher frequency of LE/Lys with a vacuolar appearance than found in control cells. Osmium and uranyl acetate stains used in electron microscopy provide contrast by staining lipid-rich membrane; the loss of electron-dense content from LE/Lys is likely due to NALL-induced lipid egress from LE/lysosomes, consistent with the reduced filipin staining shown in Figure 1. Multiple trafficking routes operate to ensure the appropriate distribution of LDL-derived cholesterol, with excess cellular cholesterol being transported to the ER for esterification by the ER-resident enzyme acyl-CoA/cholesterol acyltransferase (ACAT). We have previously identified reduced ER contact with LE/Lys in cellular models of NPC and shown that artificial expansion of ER:LE/Lys contact can rescue cholesterol accumulation in NPC (12). Following NALL treatment, LE/Lys were often in close proximity to the ER in NPC cells and ER contact with LE/Lys appeared to be increased in NALL-treated cells (Figure 2A and 2C). The abundance of membrane in areas of NPC patient fibroblasts makes it difficult to confidently distinguish ER from LE/Lys by EM. We therefore turned to a proximity ligation assay (PLA), as a fluorescent readout of membrane contact sites (25, 26). PLA using antibodies to LAMP1 on LE/Lys and VAPA on the ER confirmed the previously reported reduction in ER:LE/Lys contact in NPC cells (12). As seen by EM, ER proximity to LE/Lys was also increased in NPC patient cells treated with NALL when measured by PLA (Figure 2D and 2E, and Supplemental Figure S2A and S2B), consistent with increased ER:LE/Lysosome contact resulting from NALL treatment.

**Figure 2.**
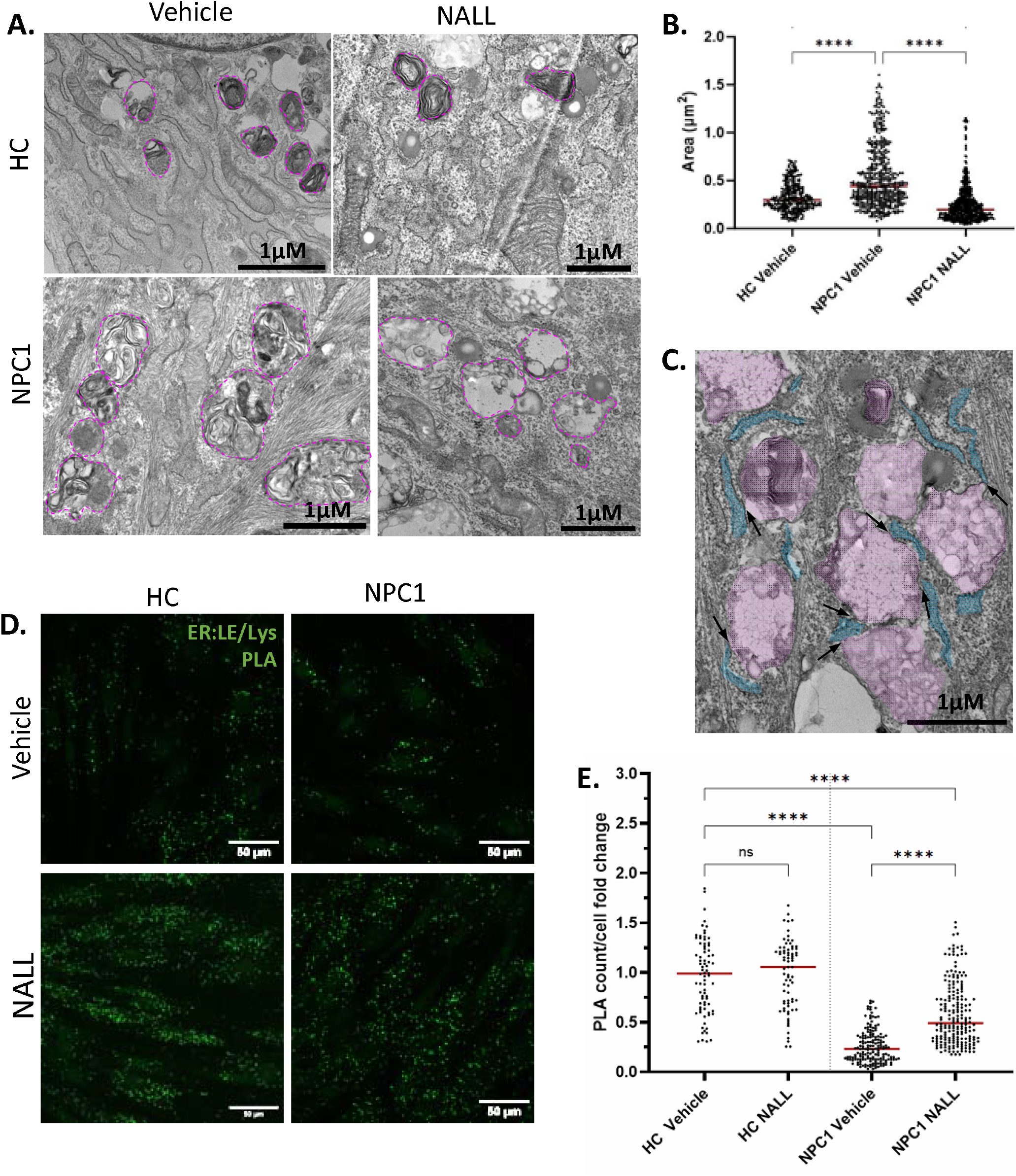
NALL restores LE/Lys morphology and ER contact sites. Primary fibroblasts from healthy control donors (HC) or NPC1 patients treated with DMSO (vehicle) or 1mM NALL for 72 hours (NALL) were fixed and prepared for electron microscopy. A) Example images showing LE/Lys (dashed magenta outline). Scale bar, 1μm. B) Area of LE/Lys per μm2 determined using Image-J. (HC: *n* = 308, NPC1: *n* = 498, NPC1 NALL: *n* = 604 cells C) Example ER:LE/Lys contact sites (black arrows) in an NPC1 patient cell treated with NALL. ER false-coloured blue and LE/Lys false-coloured pink. Scale bar, 1μm. D) Representative images of ER:LE/Lys PLA, scale bar 50μm. E) Quantitation of PLA spots/cell normalized to controls. Data analysed by one-way ANOVA followed by Tukey’s multiple comparisons test comparing all groups (* p ≤ 0.05, ** p ≤ 0.01, *** p ≤ 0.001, ns = non-significant, n = 3) (HC: *n* = 80, HC NALL: n= 75 NPC1: *n* = 174, NPC1 NALL: *n* = 204 cells).

Analysis of ER:LE/Lys contact sites in NALL-treated NPC cells by EM revealed an associated effect on lipid droplet formation. Following NALL treatment, lipid droplets were often found in close proximity to lysosomes in NPC patient cells, especially at ER:LE/Lys contact sites (Figure 3A). Indeed we found a marked increase in three-way LE/Lys:ER:lipid droplet contact sites in NALL-treated NPC cells (Figure 3A and 3B), suggesting that lipid transport from LE/Lys to the ER may promote lipid droplet formation. We therefore measured the effects of NALL on lipid droplet numbers and volume. Quantitation of the neutral lipid stain BODIPY showed a striking increase in lipid droplet formation in NPC patient cells following NALL treatment (Figure 3C and 3D and Supplemental Figure S3A and S3B). Excess cellular cholesterol is esterified by ACAT in the ER for storage in lipid droplets (27) and consistent with previous findings in CHO cells lacking NPC1 (18), NALL treatment also increased cholesterol esterification in NPC patient cells (Figure 3E). Together, these data suggest that increased ER contact likely contributes to the mechanism of NALL-mediated cholesterol clearance from LE/Lys in NPC1-deficient cells.

**Figure 3.**
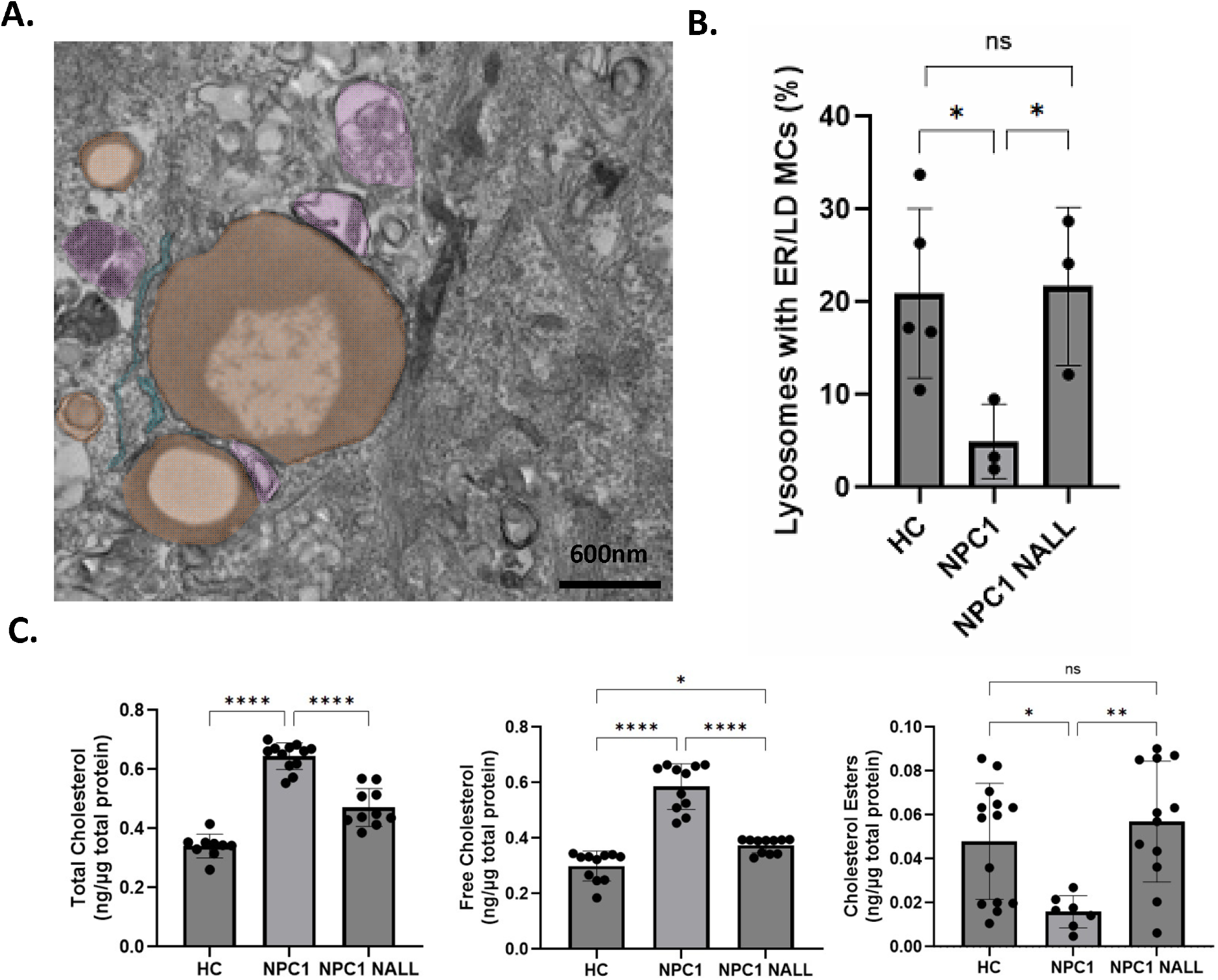
Effect of NALL on lipid droplet formation. A) Example image of LE/Lys (false-coloured pink)-ER (false-coloured blue)-lipid droplet (false-coloured orange) three-way contact sites in NPC1 patient cells treated with 1mM NALL for 72 hours. Scale bar, 600nm. B) Quantitation of the percent of LE/Lys in a 3-way contact with ER and lipid droplets. The percentage of lysosomes with an ER/LD MCS was quantified and represented as the mean of four independent experiments + SEM. Welch two-sample t-tests were performed between vehicle and treatment conditions (n=4)s. C) Amplex red cholesterol assays, expressed as cholesterol/μg of total protein. Free cholesterol (no cholesterol esterase) was subtracted from total cholesterol (with cholesterol esterase) for quantitation of cholesterol esters. One-way ANOVA was carried out and followed by Tukey’s multiple comparisons test comparing all groups C)(* p ≤ 0.05, ** p ≤ 0.01, *** p ≤ 0.001, ns = non-significant, n = 12).

### Relationship between LE/Lys cholesterol and mitochondrial contact

We and others have previously reported expansion of mitochondria:LE/Lys contact in NPC1-inhibited cells (12, 28), suggesting that increased physical connection between the two organelles could contribute to coupled lysosome and mitochondrial dysfunction in NPC, but in contrast, reduced MLCs were reported in NPC2 knockout ARPE19 cells (29). To the best of our knowledge, MLCs have not yet been examined in NPC patient cells. Consistent with previous findings, MLCs measured either by EM (Figure 4A and 4B) or by PLA using antibodies targeting LAMP1 on LE/Lys and Tom20 on mitochondria, (Figure 4C and Supplemental Figure S4A and S4B) were significantly increased in primary fibroblasts from NPC1 patients, suggesting that LE/Lys cholesterol accumulation may promote MLC formation in the absence of functional NPC1. Moreover, we found considerable variation in the number of MLCs between the three NPC1 patient cell lines (Supplemental Figure S4A and S4B), that correlates with the extent of cholesterol accumulation (Supplemental Figure S1B and S1C). To test the relationship between LDL-derived cholesterol accumulation in LE/Lys and MLC formation, we measured MLCs in cells cultured in the presence (FCS) or absence (lipoprotein-deficient serum (LPDS)) of LDL. While 24h culture in LPDS had only a minor effect on cholesterol levels, following 48h in the absence of LDL, lysosome cholesterol had normalised in NPC1 patient cells, with no significant effect on control cells (Supplemental Figure S4C an S4D). To determine the effect of reduced lysosomal cholesterol on MLC expansion in NPC, we cultured NPC1 patient cells for 48h in LPDS, and measured MLCs by PLA. Under all conditions of reduced LDL-derived cholesterol in LE/Lys, MLCs were also significantly reduced in NPC1 patient cells, consistent with high lysosomal cholesterol in NPC driving MLC expansion (Figure 4C and 4D). To further probe the effect of cholesterol clearance on MLC formation, we took advantage of our previous finding that artificial tethering of lysosomes to the ER can reverse cholesterol accumulation in cellular models of NPC (12). We first confirmed that expression of a mutant of the LE/Lys sterol-binding protein ORP1L with its sterol-binding domain deleted (ORP1L-ΔORD-GFP), expanded ER:LE/Lys contact in patient cells and reduced filipin-stained cholesterol in NPC1 patient cells (Supplemental Figure S4E and S4F), as previously reported (12). On expansion of ER contact and the associated reduction in LE/Lys cholesterol, there was a significant reduction in MLCs in NPC1 cells (Figure 4C and 4D), again consistent with high cholesterol promoting MLC expansion. Having shown that NALL treatment rescues lysosomal lipid accumulation in NPC patient cells (Figure 1A and 1B), to further probe the relationship between LE/Lys cholesterol and MLCs, we examined the effect of NALL on MLCs. Quantitation of contact sites by EM and PLA revealed that treatment with NALL reversed MLC expansion in NPC patient cells, normalising the number of contacts to that found in control cells (Figure 4C and 4D). Together these data show a striking correlation between LE/Lys cholesterol accumulation and the extent of MLC (Figure 4E).

**Figure 4.**
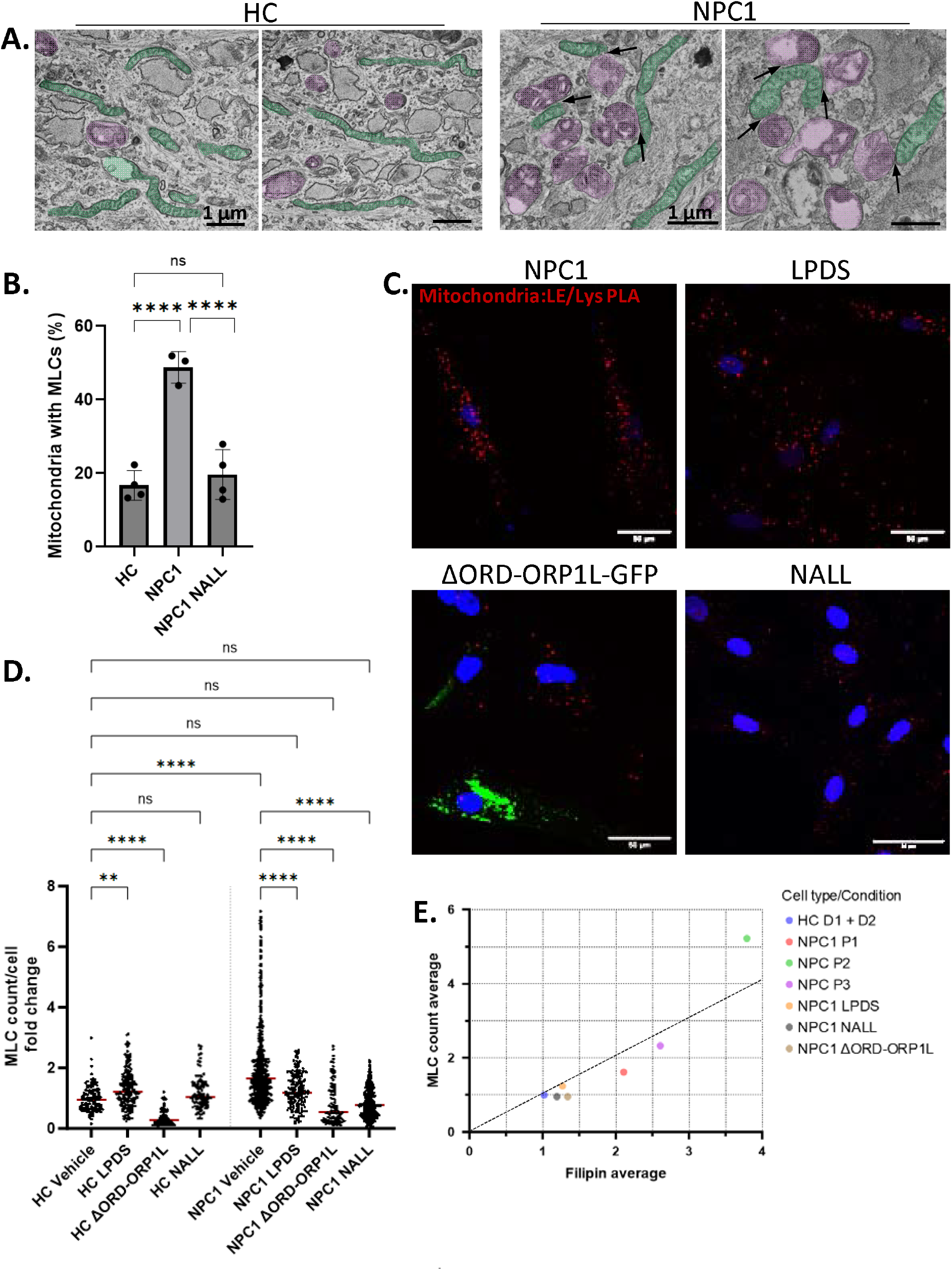
Correlation between MLCs and LE/Lys cholesterol. Primary fibroblasts from healthy control donors (HC) or NPC1 patients treated with DMSO (vehicle) or 1mM NALL for 72 hours (NALL) were fixed and prepared for electron microscopy. A) Example images showing LE/Lys false-coloured magenta and mitochondria false-coloured green. Black arrows, MLCs. Scale bar, 1μm. B) Quantitation of the percent of mitochondria with a LE/Lys contact. The percentage of mitochondria with MLCs was quantified and represented as the mean of four independent experiments + SEM. Welch two-sample t-tests were performed between vehicle and treatment conditions. C) Representative images of Mitochondria:LE/Lys PLA, scale bar 50μm. D) Quantitation of PLA spots/cell normalized to controls. One-way ANOVA was performed followed by Tukey’s multiple comparisons test comparing all groups to HC Vehicle)(* p ≤ 0.05, ** p ≤ 0.01, *** p ≤ 0.001, ns = non-significant). E) Correlation between MLCs and Filipin in NPC1 patients (P1-P3) and in cells cultured in LPDS or NALL or expressing the ER:LE/Lys tether ΔORD-ORP1L.

### Reversal of MLC expansion rescues mitochondrial dysfunction in NPC1-patient cells

As reported for other models of NPC, NALL treatment at least partially normalised the reduced basal respiration in NPC1 patient cells (Supplemental Figure S5A - S5C). Thus, treatment with NALL rescues lysosomal storage, MLC expansion and mitochondrial dysfunction in NPC models, suggesting that effects on MLCs likely contribute to the coupled restoration of LE/Lys and mitochondrial function by NALL. To further interrogate the role of MLCs in mitochondrial function, we first examined the effect of reducing LE/Lys cholesterol storage, which we found normalises MLCs in NPC patient cells (Figure 4D and 4E), on mitochondrial function. Mitochondrial membrane potential (MMP) is an indicator of mitochondrial function and has been shown to be reduced in cells lacking NPC. Using the MMP-dependent inner mitochondrial membrane marker TMRM, we also found reduced MMP in NPC1 patient fibroblasts compared with controls, that was partially restored by treatment with NALL (Figure 5A and 5B), correlating with the NALL-mediated normalisation of MLCs. Similarly, culturing cells in the absence of lipoprotein, which reduced MLCs (Figure 4E), also partially restored MMP in NPC1 patient cells (Figure 5A and 5B). We have previously shown that the LE/Lys sterol transfer proteins StARD3 is a key regulator of MLC expansion in NPC (12) and as expected, NPC1 inhibition increased MLCs in wild-type but not StARD3-knockout HeLa cells (Figure 5C and Supplemental Figure S5C). Consistent with MLC expansion contributing to mitochondrial dysfunction, the reduction in MMP on NPC1 inhibition was prevented by loss of MLC regulator StARD3 (Figure 5D and 5E), revealing an inverse correlation between MLC expansion and MMP.

**Figure 5.**
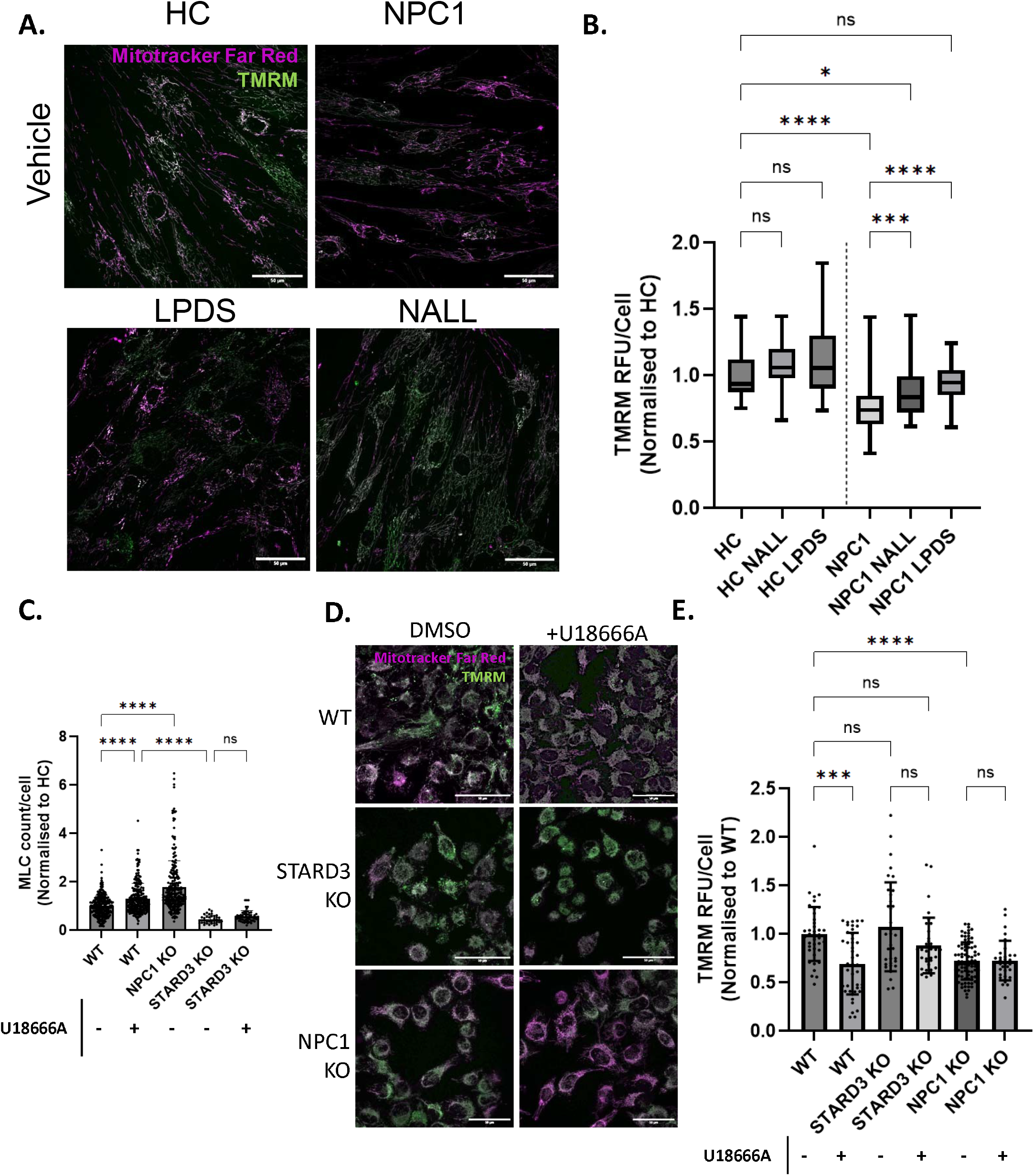
Correlation between MLCs and mitochondrial membrane potential. Primary fibroblasts from healthy control donors (HC) or NPC1 patients treated with DMSO (vehicle) or 1mM NALL for 72 hours (NALL), or LPDS for 48 hours (LPDS) were loaded with TMRM and mitotracker-647 for 30 min prior to live-cell confocal imaging. A) Representative images of TMRM (green) and mitotracker (magenta). Scale bar, 50μm. B) Quantitation of TMRM fluorescent signal (TMRM/Mitotracker normalised to HC). C) Quantitation of mitochondria:LE/Lys PLA (MLCs) in wild type (WT) or StARD3 knockout (KO) HeLa cells ± NPC1 inhibition (U18666A). D) Representative images of TMRM (green) and mitotracker (magenta) in HeLa WT or StARD3-KO cells ± NPC1 inhibition (U18666A). Scale bar, 50μm. E) Quantitation of TMRM in HeLa cells described in D, normalised to WT controls. One-way ANOVA was performed followed by Tukey’s multiple comparisons test comparing all groups to HC Vehicle)(* p ≤ 0.05, ** p ≤ 0.01, *** p ≤ 0.001, ns = non-significant).

### Restoration of lysosome repair and autophagic flux in NPC patient cells by NALL

Compromised autophagy, the pathway through which damaged organelles can be delivered to the lysosome for degradation, has been associated with cholesterol accumulation and implicated in NPC (30). A widely used marker of autophagosome formation is the lipidation of LC3-I to LC3-II (31) and an increase in LC3-II positive autophagosomes was reported in NPC models, due to impaired autophagic flux (32). We therefore tested the effect of NALL on the LC3-II by western blot. Consistent with previous reports, LC3-II was increased in NPC1 patient fibroblasts, but a 72h incubation with NALL rescued LC3-II to a level comparable with control cells (Figure 6A-6C). Bafilomycin prevents autophagosome fusion with the lysosome (33) and as expected, treatment with bafilomycin also increased LC3-II, even in the presence of NALL, indicating that a 72 hour NALL treatment does not prevent autophagosome formation (Supplemental figure S6A). These data demonstrate restoration of autophagic flux in NPC1-deficient cells by NALL treatment.

**Figure 6.**
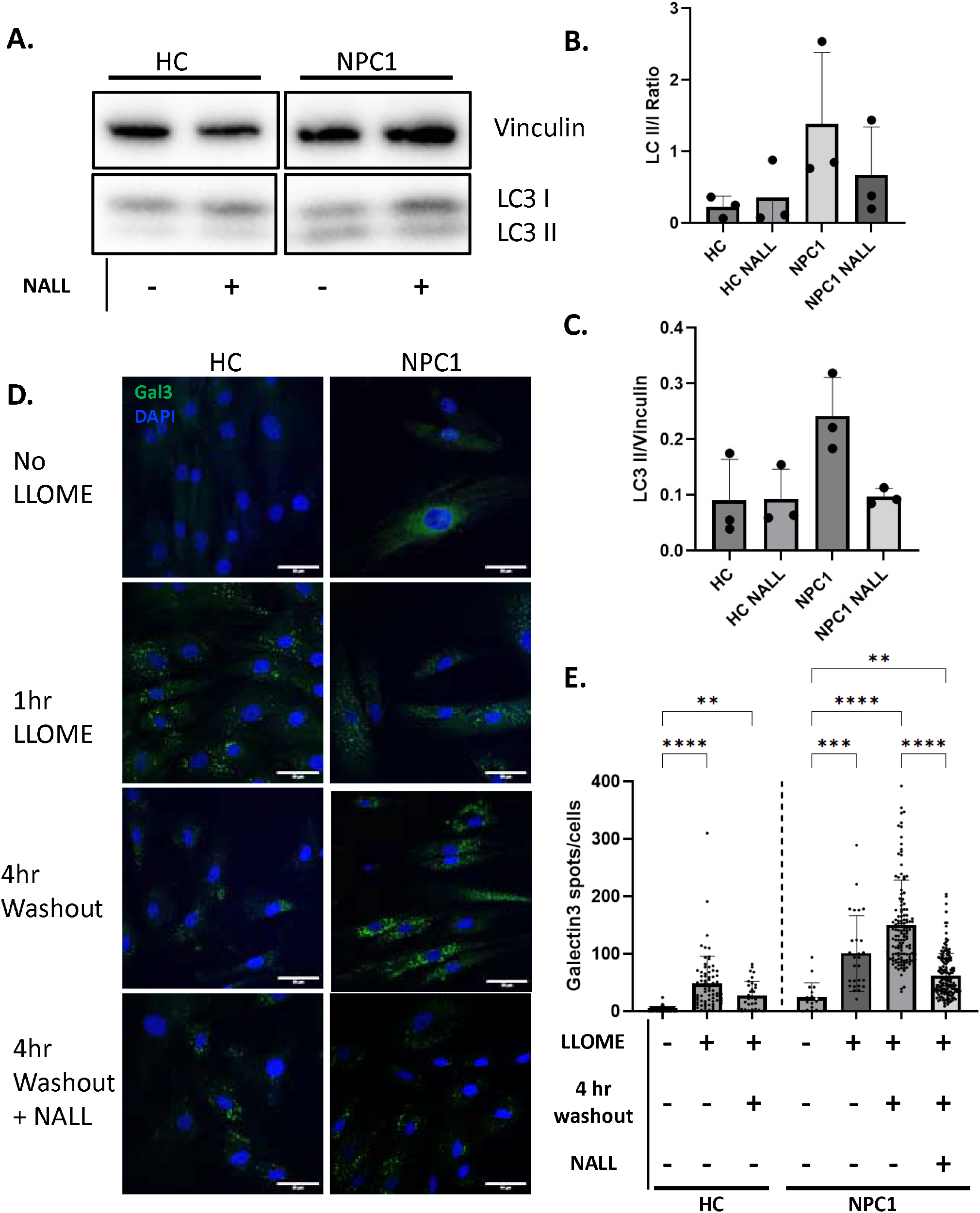
Restoration of autophagic flux and lysosome repair by NALL. A) Lysates from primary fibroblasts from healthy control donors (HC) or NPC1 patients treated with DMSO or 1mM NALL for 72 hours were immunoblotted with antibodies for the autophagy marker LC3 or loading control vinculin. B-C) quantitation of LC3-II band densitometry relative to LC3-I (B) or the vinculin loading control (C). D) Representative images of cells treated with DMSO (no LLOMe) or with 1mM LLOMe for 1 hour ± a 4 hour washout prior to fixation and staining with Galectin-3 antibody (Gal3, green) and mounting with the nuclear stain (DAPI, blue). Scale bar, 50μm. E) Quantitation of Gal3 puncta/cell shown in D. One-way ANOVA was performed followed by Tukey’s multiple comparisons test comparing all groups to HC Vehicle)(* p ≤ 0.05, ** p ≤ 0.01, *** p ≤ 0.001, ns = non-significant).

Aberrant accumulation of autophagic substrates has been proposed to contribute to lysosome damage. Galectin-3 (Gal3) is a cytosolic beta-galactoside-binding lectin, that recognises glycans exposed by LE/Lys membrane damage, to coordinate ESCRT-dependent repair and autophagic responses (34). To induce lysosome damage, cells were incubated in the presence of L-leucyl-L-leucine methyl ester (LLOMe) which is hydrolysed in the lysosome to a hydrophobic polymer widely used for its membranolytic activity (34). In both control and NPC1 patient cells, Gal3 is recruited to lysosomes following membrane damage with LLOMe, but whereas in control cells Gal3 puncta are mostly cleared after a four hour washout period, in NPC patient cells, Gal3 staining was increased after the four hour washout, indicating a gross defect in lysosome repair in NPC that was significantly improved by treatment with NALL (Figure 6D and 6E, supplemental Figure S6B).

## Discussion

Investigating the effects of NALL on coupled lysosome and mitochondrial dysfunction from an inter-organelle communication perspective, we found that NALL normalises lysosome contact with both the ER and mitochondria in NPC1 patient cells. Our demonstration that artificial expansion of ER:LE/Lys contact sites can restore cholesterol egress from LE/Lys in NPC1 patient fibroblasts (Supplemental Figure S4D and S4E) corroborates previous findings in NPC1-inhibited cells (12), offering an attractive potential mechanism for NALL-mediated cholesterol clearance. Transport of cholesterol from LE/Lys across the increased ER contact sites likely contributes to NALL-mediated reduction in LE/Lys cholesterol storage and restoration of cholesterol esterification and lipid droplet formation in NPC1 patient cells, but the increased contact is also likely to have other effects that could contribute to lipid clearance. Newly synthesized phosphatidylserine (PS) can be transported from the ER at contact sites with endosomes where it promotes recruitment of the endosomal fission protein EHD1 and therefore recycling and retrograde transport (35) which could also contribute to NALL-mediated normalisation of LE/Lys phenotypes in NPC. As well as providing sites for refilling of acidic organelle Ca^2+^ stores by the ER (25), contact sites between the ER and LE/Lys have been implicated in generating amplification of localised cytosolic Ca^2+^ signals (36, 37). Consequent activation of exocytosis (38) and autophagy (39) could contribute to both the reduced lysosome numbers and cholesterol storage that result from NALL treatment and indeed NALL treatment restored basal autophagic flux in NPC1 patient cells (Figure 6A and 6B). Interestingly, enhancement of both autophagy (24) and exocytosis (40) have proposed as potential mechanisms for the cholesterol clearance mediated by LBPA enrichment in NPC models, but rather than increasing LBPA, NALL reduced LBPA levels (Figure 1C and 1D). How that it is achieved is unclear but like cholesterol, may involve a combination of pathways including transport at contact sites, autophagy and exocytosis that together promote restoration of lysosomal homeostasis.

In a phase 3 clinical trial, a significant improvement in neurologic signs and symptoms was reported for NPC patients when receiving NALL (16), likely dependent on the restoration of both lysosomal and mitochondrial function. Mechanistically, how NALL ameliorates coupled lysosome and mitochondrial dysfunction in NPC is not understood, but extensive crosstalk exists between these two key metabolic organelles (6, 11, 14, 15). Here, we focus on direct interaction at MLCs. We found that aberrant expansion of MLCs in NPC patient cells is reversed by NALL and our data suggests that normalisation of MLCs may be key to the coupled restoration of organelle function by NALL. Taking several approaches to reducing LE/Lys cholesterol, including culturing cells in the absence of lipoprotein, we have uncovered a relationship between the cholesterol content of LE/Lys and the extent of contact with mitochondria. Our data strongly implicates the storage of LDL-derived cholesterol in LE/Lys in the formation of expanded MLCs in NPC and further indicate that these aberrant membrane contact sites contribute to the associated mitochondrial dysfunction. MLC reduction, either by cholesterol clearance from LE/Lys, or deletion of the MLC regulator StARD3, significantly improved mitochondrial membrane potential in cells lacking functional NPC1.

Together our data offer a potential mechanism for NALL therapeutic benefit: Restoration of cholesterol transport to the ER for esterification and storage in lipid droplets through pathways promoted by increased ER:LE/Lys contact lowers LE/Lys cholesterol levels in NPC1-deficient cells, reversing the pathogenic expansion of MLCs to restore mitochondrial function. These findings may pave the way for targeting membrane contact sites as a potential therapeutic strategy for other diseases.

## Methods

### Cell culture and transfection

STARD3 KO HeLa cells were kindly provided by Dr. Doris Hoglinger (University of Heidelberg) and fibroblast cells (Coriell Institute: GM5757/GM05399 Controls, GM18398/GM17921/GM22870 NPC1 patients) were cultured in DMEM F12/Glutamax/10% FBS (Invitrogen) supplemented with 10% FBS and 1% PS. Cells were transfected using Lipofectamine LTX Plus (HeLa) and nucleofection (fibroblasts) using Basic Nucleofector® Kit for Primary Mammalian Fibroblasts (Lonza), both according to manufacturer’s instructions.

### Antibodies

Mouse anti-TOM20 (D8T4N, diluted 1:400) was purchased from Santa Cruz Biotechnology, rabbit anti-LAMP1 (D2D11, diluted 1:500) and rabbit anti-LC3 (diluted 1:1000) were from Cell Signaling Technology, and rabbit anti-Vinculin (ab91459, diluted 1:10,000) was from Abcam. Mouse anti-LBPA (clone 6C4, diluted 1:200) was purchased from Echelon Biosciences, and rat anti-mouse/human Mac-2 (Galectin-3, clone M3/38, diluted 1:200) was from BioLegend. TMRM was used at a final concentration of 20 nM, MitoTracker at 500 nM, and LysoTracker Red at 1 μM. Alexa Fluor Plus 750 Phalloidin was used at a dilution of 1:1000.

### Western blotting and immunoprecipitation

For Western blotting, cells were lysed in lysis buffer [40 mM HEPES, 80 mM NaCl, 10 mM EDTA, 10 mM EGTA, 1% Triton-X100, protease inhibitor cocktail (Calbiochem set I), phosphatase inhibitor cocktail (Calbiochem set II)]. Lysates were fractioned by SDS–PAGE on 10% gels under reducing conditions and immunoblotted on PVDF membranes. Membranes were incubated overnight at 4°c with primary antibodies (1:1000 LC3, 1:10000 Vinculin) Following incubation with HRP-conjugated secondary antibodies (1:1000), membranes were scanned in BioRad ChemiDoc imaging system.

### Filipin Staining

Cells were fixed in 4% PFA for 20 min, quenched with 20 mM glycine, and stained with filipin (100 µg/mL) in PBS for 30–60 min at RT in the dark. Imaging was performed using a 405 nm laser line with emission collection up to 500 nm.

### Proximity Ligation Assay (PLA)

Cells grown on 10-mm coverslips were fixed in 4% paraformaldehyde (PFA) for 20 min at room temperature (RT), quenched in 15 mM glycine in PBS for 10 min, and permeabilized with 0.1% Triton X-100 in PBS for 10 min at RT. After two PBS washes, blocking was performed in 1% bovine serum albumin (BSA) in PBS for 1 hour at RT. For ER:LE/Lys contact site analysis, primary antibodies against LAMP1 (rabbit anti-LAMP1, Cell Signaling, #9091; 1:160) and VAPA (???) were used; for MLC analysis, antibodies were targeting LAMP1 (rabbit anti-LAMP1, Cell Signaling, #9091; 1:160) and TOM20 (mouse anti-TOM20, Santa Cruz, sc-17764; 1:100). Antibodies were diluted in 1% BSA and incubated overnight at 4 °C. Coverslips were washed twice in Wash Buffer A (Duolink kit, Sigma, DUO92101) and incubated with Duolink anti-rabbit PLUS and anti-mouse MINUS PLA probes (1:10 dilution) at 37°C for 1 h in a humidity chamber. After washing, ligation was performed at 37°C for 30 min, followed by amplification at 37°C for 100 min, protected from light. Coverslips were washed in Wash Buffer B, briefly rinsed in 0.01× Wash Buffer B, and mounted with Duolink mounting media.

Images were acquired using a Leica Stellaris 8 confocal microscope under consistent exposure settings. PLA puncta were quantified in ImageJ.

### Mitochondrial Respiration Analysis Using Seahorse XF

Cellular respiration was assessed using the Seahorse XF Pro Analyzer. Cells (5,000 cells/well) were seeded into XF96 microplates two days prior to assay. On the day of the experiment, cells were equilibrated in unbuffered DMEM (25 mM glucose, 2 mM L-glutamine, 1 mM pyruvate, pH 7.4) in a non-CO_2_ incubator at 37°C for 1 h. Sensor cartridges were prehydrated overnight in XF Calibrant at 37°C.

Mitochondrial stress test compounds—oligomycin (1 µM), FCCP (2 µM), and rotenone/antimycin A (0.5 µM each)—were sequentially injected. Oxygen consumption rate (OCR) measurements were recorded automatically. Following the assay, nuclei were stained with DAPI and counted using a BioTek Cytation 10 for normalization.

Mitochondrial parameters were calculated as per Seahorse Analytics guidelines and statistical analyses were performed using GraphPad Prism.

### Mitochondrial Membrane Potential - TMRM

Before imaging, the fluorodishes were washed with a recording buffer solution DMEM/F-12, followed by incubating the cells in recording buffer solution containing 20 nM TMRM (Thermo Fisher Scientific, T668) and MitoTracker Deep Red FM (500nm) for 30 minutes at 37°C. The fluorodishes were washed with the recording buffer and fresh buffer solution was added containing 20 nM TMRM. Images were acquired using a Leica Stellaris 8 confocal microscope under consistent exposure settings in 37°C 5% CO2 environment chamber. Z-stacks were acquired for TMRM and Mitotracker. TMRM fluorescence was excited at 543 nm with emission maxima at 574 nm. The obtained images were later processed using Fiji/ImageJ.

### Cholesterol Quantification by Amplex Red Assay

Intracellular cholesterol was quantified using the Amplex™ Red Cholesterol Assay Kit (Invitrogen, A12216) according to the manufacturer’s instructions, adapted from Gong et al., 2019.

Briefly, cells were grown in 6-well plates and cell lysates were harvested in assay buffer, water sonicated for 30 seconds and diluted to fall within the linear detection range. Reactions were set up in 96-well black plates containing 50 μL of sample or standard, 50 μL of working solution (containing 300 μM Amplex Red reagent, 2 U/mL HRP, 2 U/mL cholesterol oxidase, +/-0.2 U/mL cholesterol esterase in order to measure total and unesterified cholesterol).

The fluorescence (excitation 540 nm, emission 590 nm) was measured after 30 min incubation at 37°C in the dark. Total cholesterol was normalised to protein concentration determined by BCA assay.

### Transmission Electron Microscopy (TEM)

Cells were washed in PBS and fixed in 2% PFA/2% glutaraldehyde in 0.1 M cacodylate buffer (pH 7.4) for 30 min at RT. After rinsing in 0.1 M and then 0.05 M cacodylate, samples were post-fixed with 1% osmium tetroxide/1.5% potassium ferricyanide in 0.1 M cacodylate buffer for 1 hour at 4°C. Cells were stained en bloc with 2% uranyl acetate replacement solution (UA Zero) for 1 hour at RT, dehydrated through graded ethanol series, and infiltrated with TAAB 812 epoxy resin (TAAB Laboratories). Polymerization was performed overnight at 60°C.

Ultrathin sections (70 nm) were cut and imaged on a JEOL 1400Plus EM (JEOL ltd, Tokyo, Japan) fitted with an Advanced Microscopy Technologies (AMT) NanoSprint12 camera (AMT Imaging Direct, Woburn, MA, USA).

## Author contributions

E.E., S.K. and F.P. conceived the project. S.K conducted all fluorescence imaging, electron microscopy, Amplex Red assays and western blotting with help from A.M and did all of the data analysis and figure preparation. E.E. wrote the first draft, F.P and S.K reviewed and edited.

## Acknowledgements

This work was supported by the Medical Research Council research grant (grant no. MR/V013882/1) and DTP (grant no. MR/N01386711). We are grateful to Doris Höglinger (Heidelberg University, 69120 Heidelberg, Germany) for cells and Jacques Neefjes (Leiden University Medical Center, Netherlands) for the ORP1L-deltaORD-GFP plasmid.

