## Supplemental Figures for "N-Acetyl-l-Leucine (NALL) rescues inter-organelle communication in Niemann-Pick disease type-C patient cells"

### Slide 1
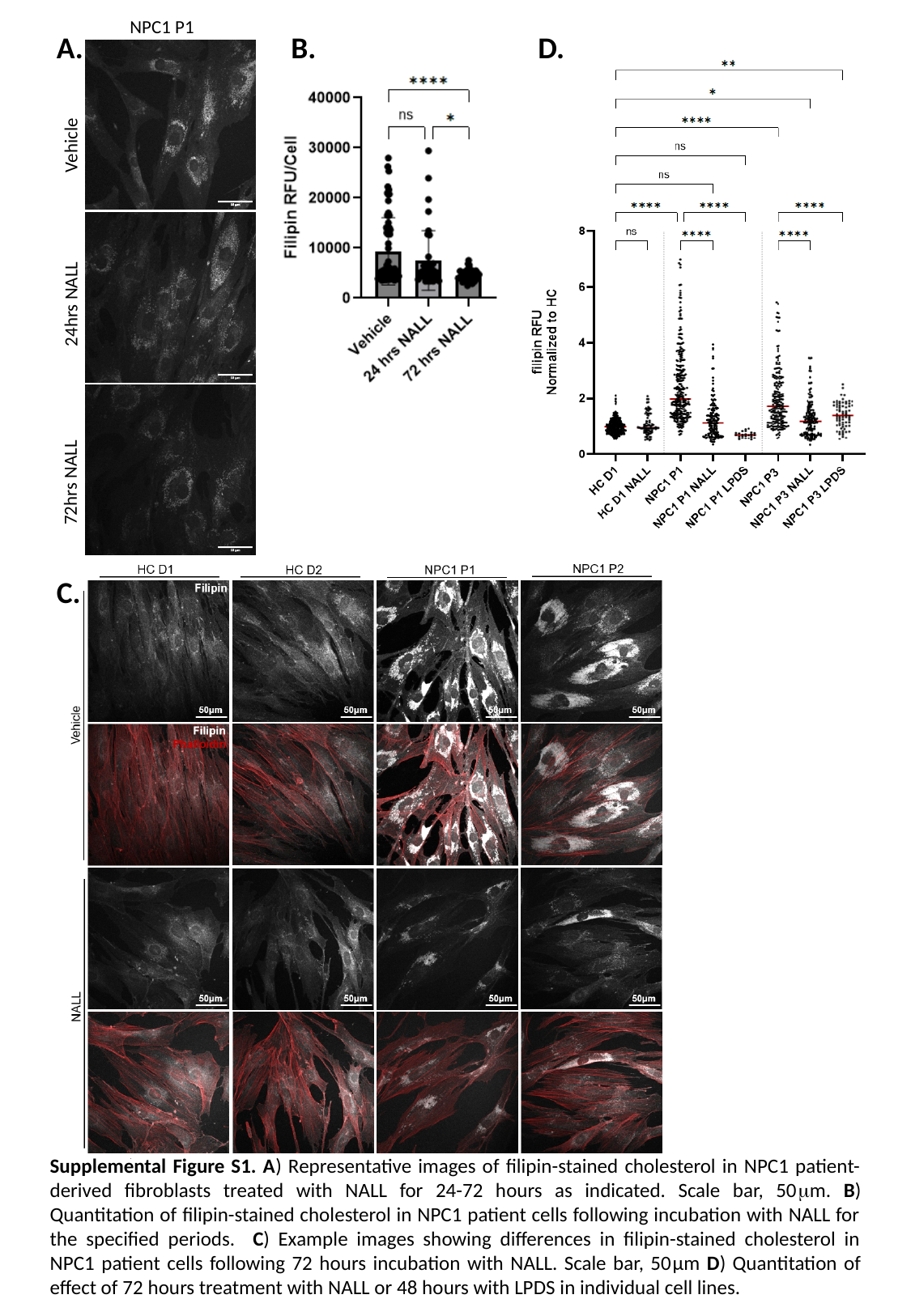

NPC1 P1
A.
B.
D.
Vehicle
24hrs NALL
72hrs NALL
C.
Supplemental Figure S1. A) Representative images of filipin-stained cholesterol in NPC1 patient-derived fibroblasts treated with NALL for 24-72 hours as indicated. Scale bar, 50μm. B) Quantitation of filipin-stained cholesterol in NPC1 patient cells following incubation with NALL for the specified periods. C) Example images showing differences in filipin-stained cholesterol in NPC1 patient cells following 72 hours incubation with NALL. Scale bar, 50μm D) Quantitation of effect of 72 hours treatment with NALL or 48 hours with LPDS in individual cell lines.

### Slide 2
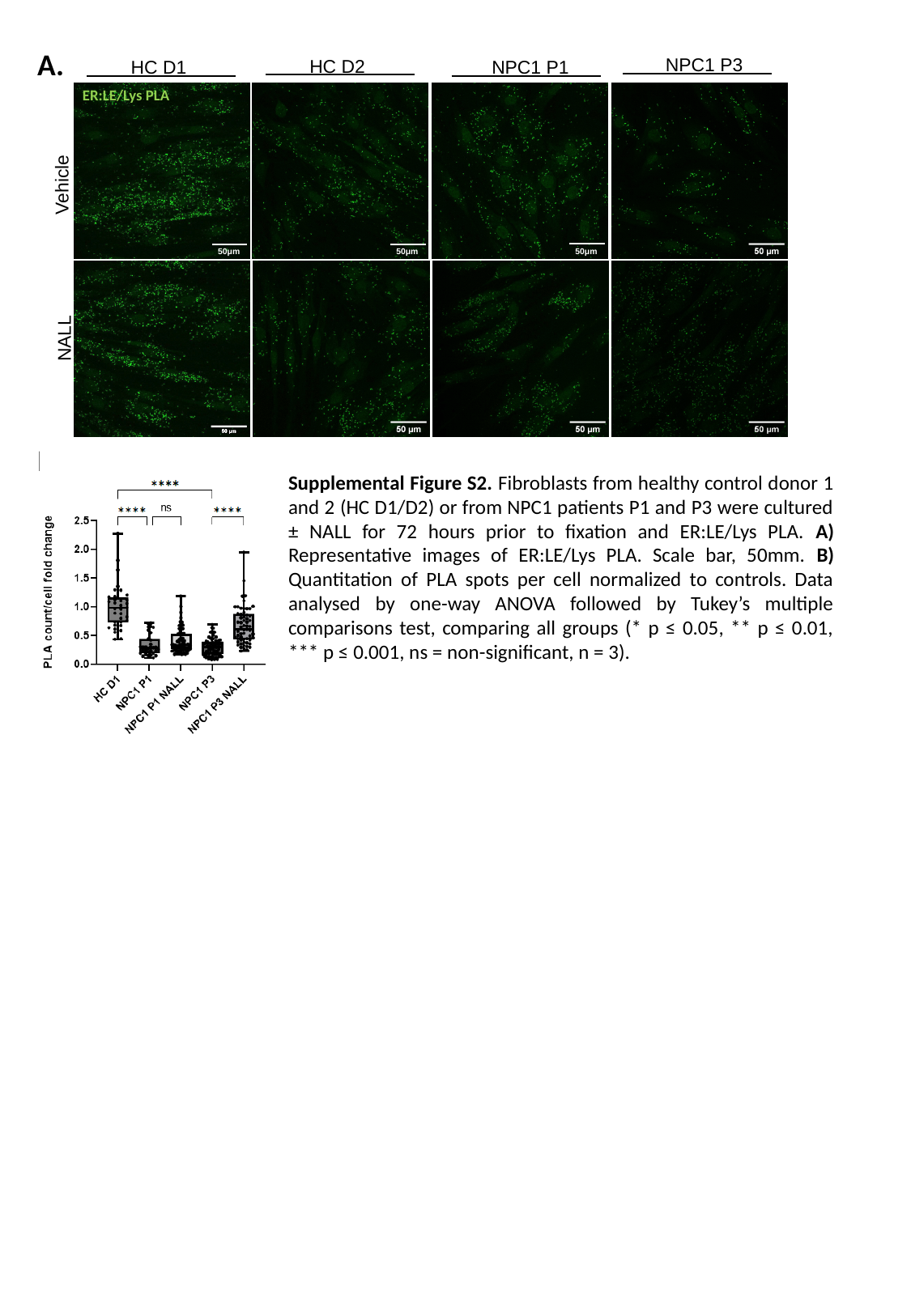

A.
NPC1 P3
HC D2
HC D1
NPC1 P1
Vehicle
NALL
ER:LE/Lys PLA
50μm
50μm
50μm
B.
Supplemental Figure S2. Fibroblasts from healthy control donor 1 and 2 (HC D1/D2) or from NPC1 patients P1 and P3 were cultured ± NALL for 72 hours prior to fixation and ER:LE/Lys PLA. A) Representative images of ER:LE/Lys PLA. Scale bar, 50mm. B) Quantitation of PLA spots per cell normalized to controls. Data analysed by one-way ANOVA followed by Tukey’s multiple comparisons test, comparing all groups (* p ≤ 0.05, ** p ≤ 0.01, *** p ≤ 0.001, ns = non-significant, n = 3).
50μm

### Slide 3
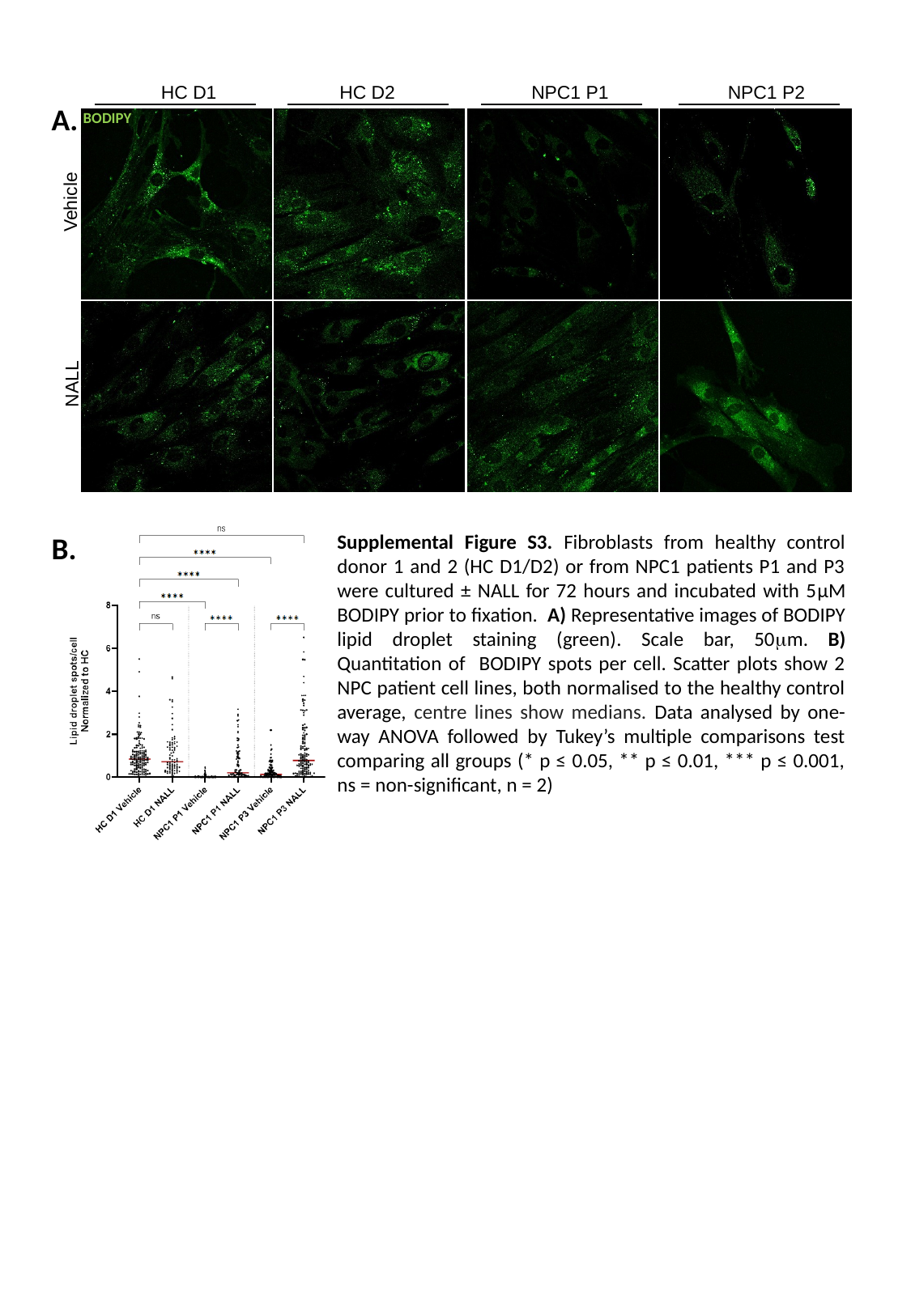

HC D1
HC D2
NPC1 P1
NPC1 P2
Vehicle
NALL
A.
BODIPY
B.
Supplemental Figure S3. Fibroblasts from healthy control donor 1 and 2 (HC D1/D2) or from NPC1 patients P1 and P3 were cultured ± NALL for 72 hours and incubated with 5μM BODIPY prior to fixation. A) Representative images of BODIPY lipid droplet staining (green). Scale bar, 50μm. B) Quantitation of BODIPY spots per cell. Scatter plots show 2 NPC patient cell lines, both normalised to the healthy control average, centre lines show medians. Data analysed by one-way ANOVA followed by Tukey’s multiple comparisons test comparing all groups (* p ≤ 0.05, ** p ≤ 0.01, *** p ≤ 0.001, ns = non-significant, n = 2)

### Slide 4
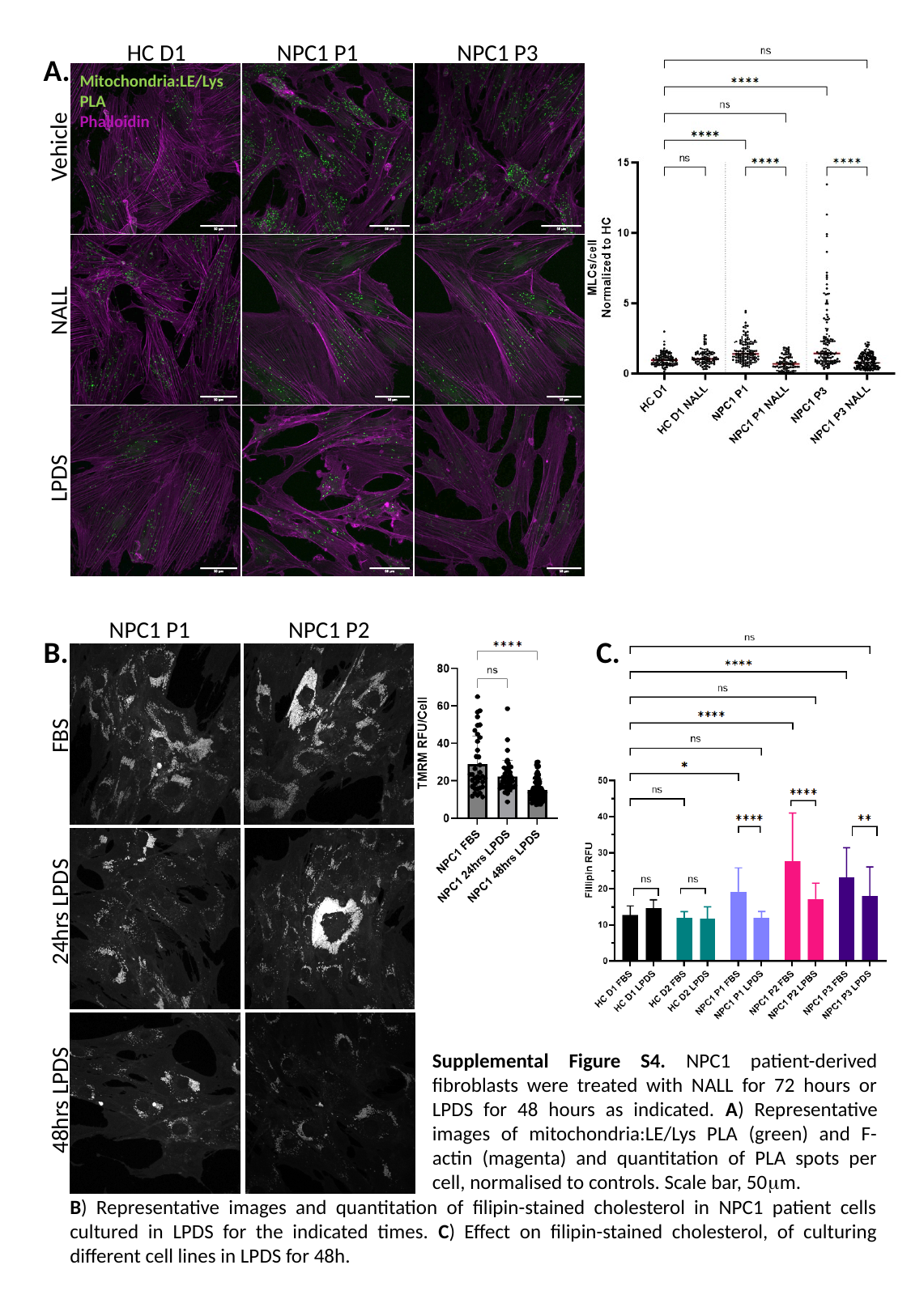

HC D1
NPC1 P1
NPC1 P3
Mitochondria:LE/Lys PLA
Phalloidin
Vehicle
NALL
LPDS
A.
NPC1 P1
NPC1 P2
FBS
24hrs LPDS
48hrs LPDS
B.
C.
Supplemental Figure S4. NPC1 patient-derived fibroblasts were treated with NALL for 72 hours or LPDS for 48 hours as indicated. A) Representative images of mitochondria:LE/Lys PLA (green) and F-actin (magenta) and quantitation of PLA spots per cell, normalised to controls. Scale bar, 50μm.
B) Representative images and quantitation of filipin-stained cholesterol in NPC1 patient cells cultured in LPDS for the indicated times. C) Effect on filipin-stained cholesterol, of culturing different cell lines in LPDS for 48h.

### Slide 5
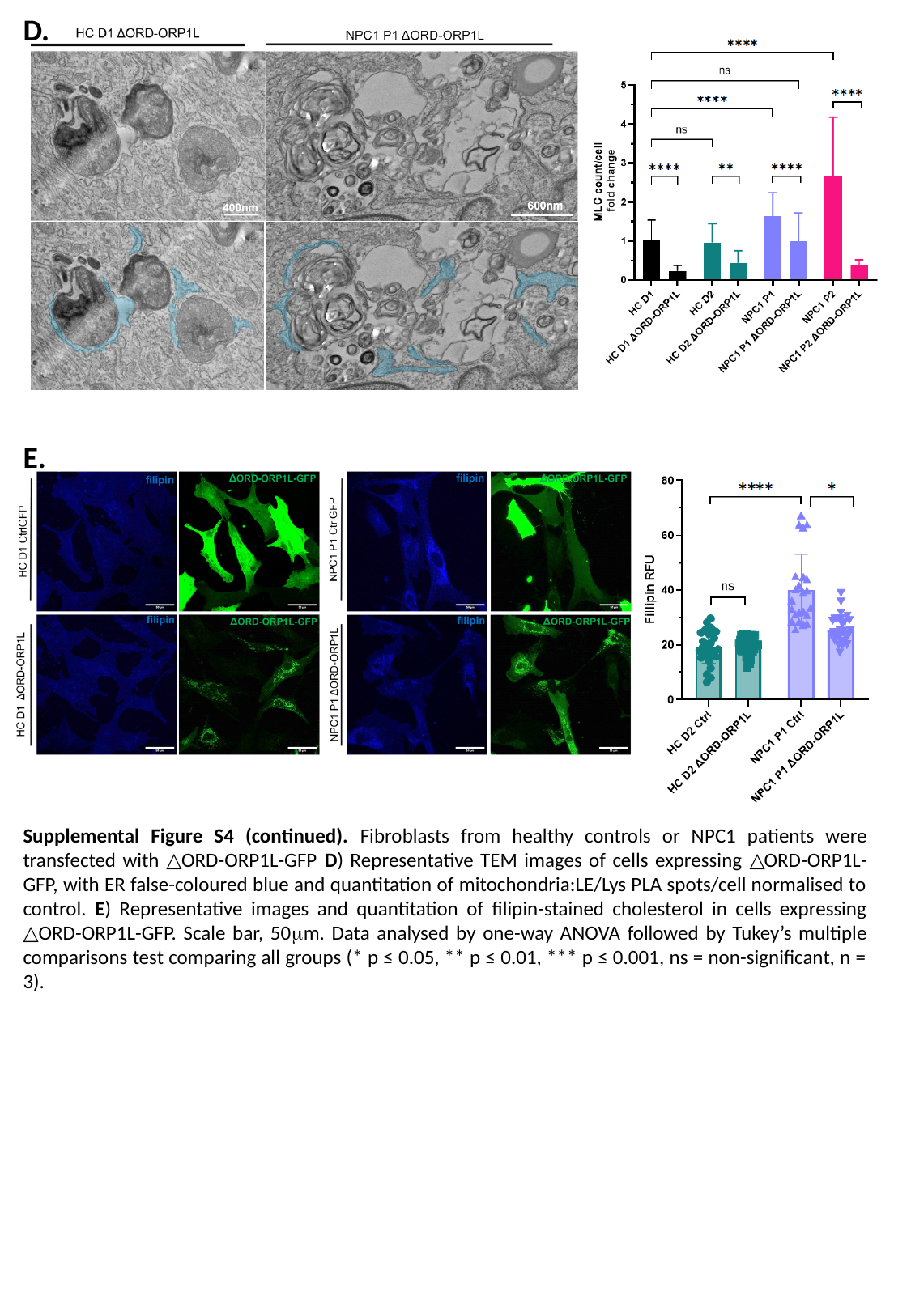

D.
E.
Supplemental Figure S4 (continued). Fibroblasts from healthy controls or NPC1 patients were transfected with △ORD-ORP1L-GFP D) Representative TEM images of cells expressing △ORD-ORP1L-GFP, with ER false-coloured blue and quantitation of mitochondria:LE/Lys PLA spots/cell normalised to control. E) Representative images and quantitation of filipin-stained cholesterol in cells expressing △ORD-ORP1L-GFP. Scale bar, 50μm. Data analysed by one-way ANOVA followed by Tukey’s multiple comparisons test comparing all groups (* p ≤ 0.05, ** p ≤ 0.01, *** p ≤ 0.001, ns = non-significant, n = 3).

### Slide 6
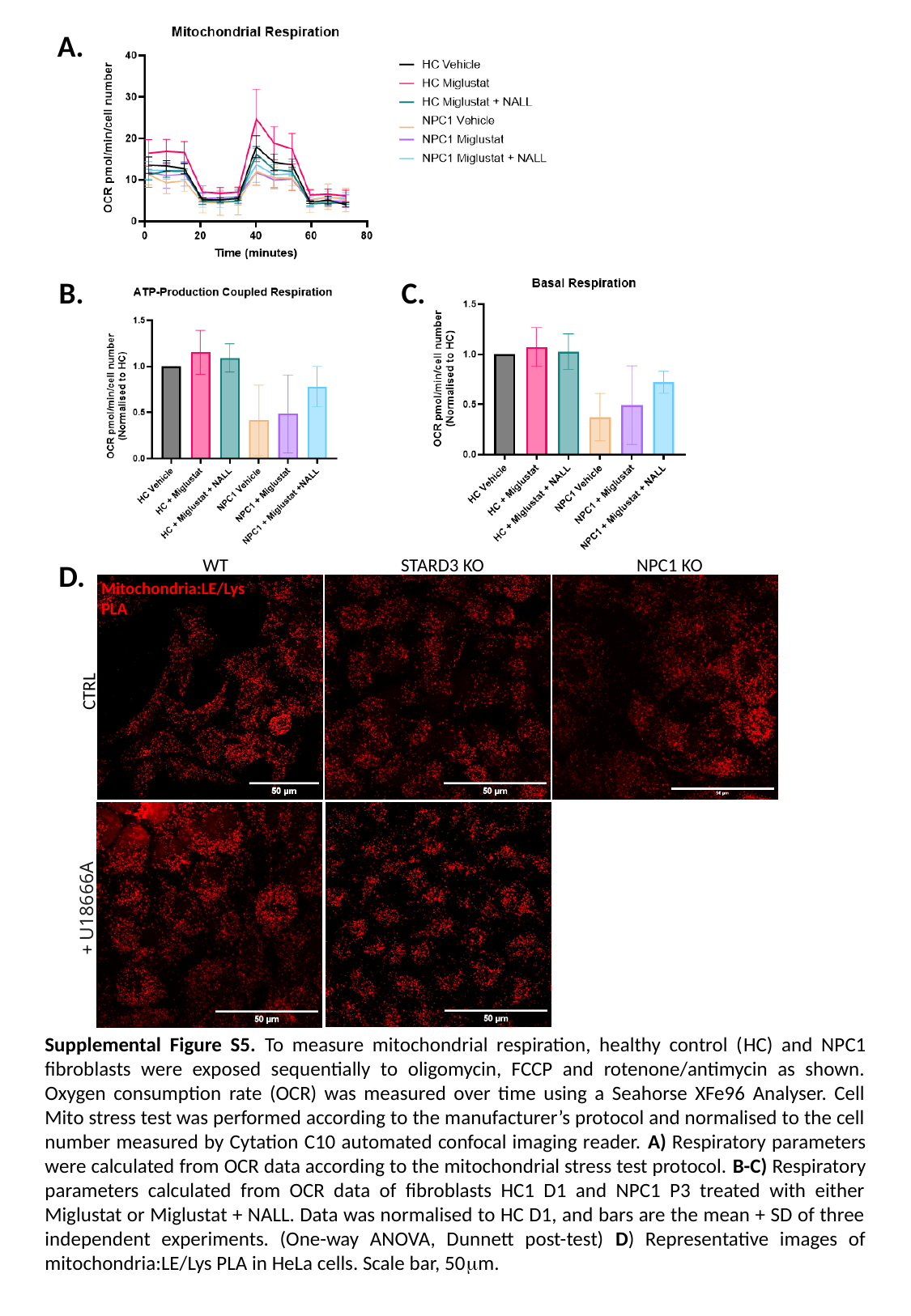

A.
B.
C.
WT
STARD3 KO
NPC1 KO
CTRL
+ U18666A
D.
Mitochondria:LE/Lys PLA
Supplemental Figure S5. To measure mitochondrial respiration, healthy control (HC) and NPC1 fibroblasts were exposed sequentially to oligomycin, FCCP and rotenone/antimycin as shown. Oxygen consumption rate (OCR) was measured over time using a Seahorse XFe96 Analyser. Cell Mito stress test was performed according to the manufacturer’s protocol and normalised to the cell number measured by Cytation C10 automated confocal imaging reader. A) Respiratory parameters were calculated from OCR data according to the mitochondrial stress test protocol. B-C) Respiratory parameters calculated from OCR data of fibroblasts HC1 D1 and NPC1 P3 treated with either Miglustat or Miglustat + NALL. Data was normalised to HC D1, and bars are the mean + SD of three independent experiments. (One-way ANOVA, Dunnett post-test) D) Representative images of mitochondria:LE/Lys PLA in HeLa cells. Scale bar, 50μm.

### Slide 7
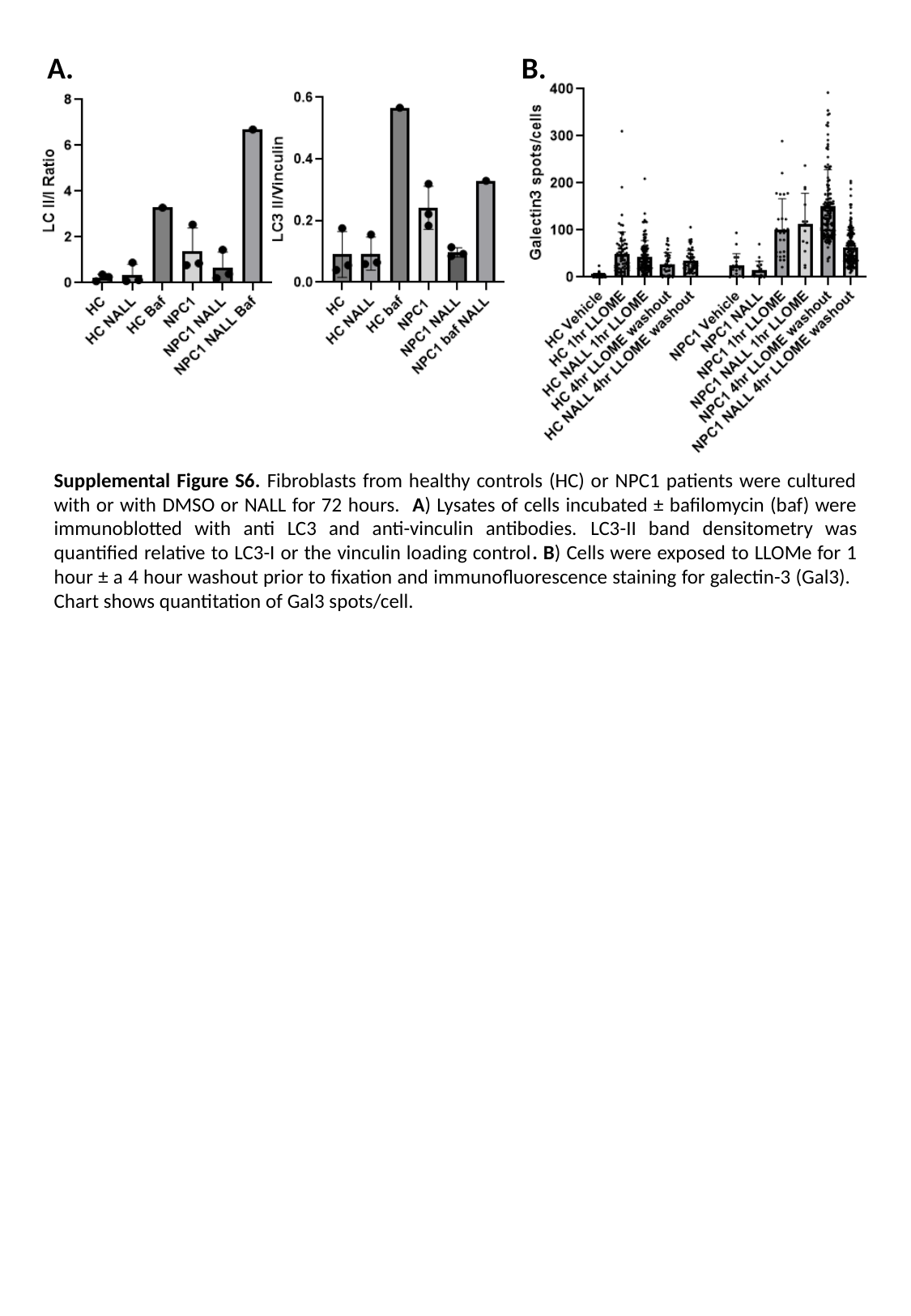

A.
B.
Supplemental Figure S6. Fibroblasts from healthy controls (HC) or NPC1 patients were cultured with or with DMSO or NALL for 72 hours. A) Lysates of cells incubated ± bafilomycin (baf) were immunoblotted with anti LC3 and anti-vinculin antibodies. LC3-II band densitometry was quantified relative to LC3-I or the vinculin loading control. B) Cells were exposed to LLOMe for 1 hour ± a 4 hour washout prior to fixation and immunofluorescence staining for galectin-3 (Gal3). Chart shows quantitation of Gal3 spots/cell.
